# Differential reorganization of episodic and semantic memory systems in epilepsy-related mesiotemporal pathology

**DOI:** 10.1101/2023.09.28.560002

**Authors:** Donna Gift Cabalo, Jordan DeKraker, Jessica Royer, Ke Xie, Shahin Tavakol, Raúl Rodríguez-Cruces, Andrea Bernasconi, Neda Bernasconi, Alexander Weil, Raluca Pana, Birgit Frauscher, Lorenzo Caciagli, Elizabeth Jefferies, Jonathan Smallwood, Boris C. Bernhardt

**Affiliations:** Multimodal Imaging and Connectome Analysis Laboratory, McConnell Brain Imaging Centre, Montreal Neurological Institute, McGill University, Montreal, QC, Canada; Montreal Neurological Institute and Hospital, McGill University, Montreal, QC, Canada; Analytical Neurophysiology Laboratory, Montreal Neurological Institute, McGill University, Montreal, Quebec, Canada; Neuroimaging of Epilepsy Laboratory, McConnell Brain Imaging Centre, Montreal Neurological Institute and Hospital, McGill University, Montreal, QC, Canada; CHU St Justine, Montreal, QC; Department of Bioengineering, University of Pennsylvania, Philadelphia, USA; University of York, North Yorkshire, United Kingdom; Queen’s University, Kingston, Ontario, Canada

**Keywords:** declarative memory, mesiotemporal lobe, neuroimaging, connectomics, gradients, epilepsy

## Abstract

Declarative memory encompasses episodic and semantic divisions. Episodic memory captures singular events with specific spatiotemporal relationships, while semantic memory houses context-independent knowledge. Behavioural and functional neuroimaging studies have revealed common and distinct neural substrates of both memory systems, implicating mesiotemporal lobe (MTL) regions such as the hippocampus and distributed neocortices. Here, we explored declarative memory system reorganization in patients with unilateral temporal lobe epilepsy (TLE), as a human disease model to test the impact of variable degrees of MTL pathology on memory function. Our cohort included 31 patients with TLE as well as 60 age and sex-matched healthy controls and all participants underwent episodic and semantic retrieval tasks during a multimodal MRI session. The functional MRI tasks were closely matched in terms of stimuli and trial design. Capitalizing on non-linear connectome gradient mapping techniques, we derived task-based functional topographies during episodic and semantic memory states, both in the MTL and in neocortical networks. Comparing neocortical and hippocampal functional gradients between TLE patients and healthy controls, we observed a marked topographic reorganization of both neocortical and MTL systems during episodic memory states. In the neocortex, alterations were characterized by reduced functional differentiation in patients in lateral temporal and midline parietal cortices in both hemispheres. In the MTL, on the other hand, patients presented with a more marked functional differentiation of posterior and anterior hippocampal segments ipsilateral to the seizure focus and pathological core, indicating perturbed intrahippocampal connectivity. Semantic memory reorganization was also found in bilateral lateral temporal and ipsilateral angular regions, while hippocampal functional topographies were unaffected. Leveraging MRI proxies of MTL pathology, we furthermore observed alterations in hippocampal microstructure and morphology are associated with TLE-related functional reorganization during episodic memory. Moreover, correlation analysis and statistical mediation models revealed that these functional alterations contributed to behavioural deficits in episodic, but again not semantic memory in patients. Altogether, our findings suggest that semantic processes rely on distributed neocortical networks, while episodic processes are supported by a network involving both the hippocampus and neocortex. Alterations of such networks can provide a compact signature of state-dependent reorganization in conditions associated with MTL damage, such as TLE.

## Introduction

Declarative memory is commonly divided into episodic and semantic memory components. Episodic memories are unique events with specific spatiotemporal associations, while semantic memory refers to context-invariant knowledge and facts about the world.^1,2,3^ The relationship between both forms of memory is complex, with behavioural and neuroimaging studies suggesting shared and distinct substrates. Behavioural studies have shown a synergistic interplay between both memory types, where episodic memory for events and words processed with semantic categorization, exceeds those from non-semantic tasks.^4,5,6^ Functional and structural neuroimaging evidence in healthy populations supports this finding, suggesting shared neural substrates across both memory types. Shared key networks are situated in the medial and lateral temporal lobes,^7,8, 9,10,11,12,13–16,17,18^ alongside posterior parietal regions.^19–21^ More broadly, networks involved in both memory types have been shown to engage similar intrinsic functional systems, notably the fronto-parietal network^22–24^ (FPN) and the default mode network (DMN).^10,25,26^ As such, behavioural and neural evidence collectively implicates shared processes in both forms of declarative memory. On the other hand, findings also point to some divergences.

Generally, episodic processes appear to rely more on medial temporal lobe (MTL) systems, while anterior temporal lobe (ATL) regions are more heavily implicated in semantic processing.^10,16^ Similarly, prior work has suggested differential activations in middle frontal *vs* inferior temporal regions in episodic as compared to semantic retrieval. ^27^ A neural divergence between both memory systems is supported by the *(1)* controlled semantic cognition framework, which stipulates that semantic representations within the ATL ‘hub’ and modality-specific ‘spokes’ interact with a ‘semantic’ control’ system that dynamically supervises and adjusts semantic retrieval bilaterally,^1,28–30^ and *(2)* hippocampus-neocortical connectivity, such that the hippocampus interacts closely with other regions through process-specific alliances to support episodic memory.^13,31,32^ A systematic investigation of neocortical and MTL networks will help to understand substrates contributing to declarative memory function in the human brain.

Temporal lobe epilepsy (TLE) is a common pharmaco-resistant epilepsy in adults and can serve as a human disease model to probe declarative memory system reorganization in young and middle-aged adults.^32,33^ Patients typically present with pathology of structures in the MTL, notably the hippocampus^34–37^ and prior research has demonstrated marked impairments in episodic memory, in some patients even at a young age.^32,38–40^ Episodic memory impairment has been directly related to degrees of hippocampal structural alterations,^41–46^ together with widespread structural and functional imbalances in both temporal and extratemporal neocortical networks.^47,48^ In contrast to the relatively consistent finding of episodic memory impairment in TLE, semantic memory has been less frequently studied, and reports remain mixed. Semantic memory function appears to be rather mildly affected,^46,49^ even in TLE patients with pronounced MTL hypometabolism.^49^ Moreover, and in contrast to the frequent finding of episodic memory disruptions in patients undergoing selective MTL resection^50–52^ there are less marked semantic memory impairments seen postsurgically.^53,54^ Such resilience suggests a more distributed cortical control network substrate for semantic memory involving both hemispheres,^55,56^ in contrast to MTL representations of episodic memory.^57,58^ Individuals with TLE may, therefore, have difficulty recalling specific events or episodes from their past while retaining their general knowledge of the world. However, other studies have found significant semantic memory deficits in patients with TLE,^59–61^ which have been associated with MTL^41^ and ATL lesions.^62^ Additionally, poor performance on semantic memory tasks has been demonstrated in TLE patients after unilateral ATL resection.^59,63,64^ Nevertheless, as impairments in declarative memory challenge the day-to-day functioning and well-being of patients, it is critical to derive more mechanistic insights into these memory types and their differential vulnerabilities in TLE.

Contemporary systems neuroscience has emphasized a key role of the spatial organization of macroscale functional networks in human cognition.^65–68^ Recent analytical and conceptual advances have notably emphasized the existence and utility of spatial *gradients*^65,68–71^ as low-dimensional axes of cortical organization, often reflective of established principles of cortical topography and hierarchy.^65,72,73^ At the level of intrinsic functional connectivity derived from task-free functional MRI, the first/principal gradient anchors primary sensory and motor areas at one end and transmodal association systems, like the FPN and DMN, at the other end, whereas a second gradient differentiates visual and somatosensory/motor areas.^65^ Gradients also compactly describe the subregional hippocampal organization, with a first gradient differentiating anterior-posterior (long axis) divisions and a second gradient the proximal-distal arrangement of different subfields along the infolding of this allocortical structure.^74–77^ Beyond the application of gradient mapping techniques to study healthy brain organization,^65,76,78–85^ recent studies have interrogated gradient alterations in diseased cohorts, including TLE patients.^86,87^ A recent study from our group demonstrated microstructural gradient contractions in TLE in the neocortex, which were primarily localized in temporo-limbic systems and found to track deficits in episodic memory recall accuracy.^87^ Another study demonstrated functional gradient expansion in subcortical structures during resting conditions, with notable differences in ipsilateral hippocampal gradient deviations between left *vs* right-lateralized TLE.^86^ Additionally, patterns of cortical asymmetry and atrophy were shown to have been distinctively linked with microstructural and functional gradients.^88^ Gradient mapping, therefore, offers a framework to interrogate global memory network topographies and their association with cognition in both health and disease.

The current study investigated declarative memory network organization in healthy individuals and a cohort of TLE patients presenting with variable degrees of MTL pathology. All participants underwent episodic and semantic retrieval tasks inside the MRI scanner, with both tasks being closely matched for stimulus presentation, task design, and task structure.^32,46^ For each of the declarative memory states, we mapped spatial gradients of memory state connectivity, both within the MTL and in macroscale neocortical networks. By adopting gradients as an analytical and conceptual framework for task-derived functional MRI data in TLE, we were able to answer the following questions: (*i*) whether networks subserving episodic *vs* semantic memory in patients undergo a shared or selective reorganization relative to controls, *(ii)* whether memory network reorganization reflects TLE-related structural alterations in the MTL, capitalizing on established *in vivo* proxies of mesiotemporal pathology^35,89^ ^90,91^ *(iii)* whether these findings reflect behavioural impairment in relational memory seen in patients.

## Materials and methods

### Participants

We studied 31 patients with drug-resistant TLE (17 females, age=35±11.59 years, left/right handed=1/30), and 60 healthy controls (30 males, age=33± 8 years, left/right handed=2/58). All participants were recruited between May 2018 and March 2023 at the Montreal Neurological Institute (MNI). Controls met the following inclusion criteria: (1) age between 18-65 years; (2) no neurological or psychiatric illness; (3) no MRI contraindication; (4) no drug/alcohol abuse problem; and (5) no history of brain injury and surgery. Patient demographics and clinical features were obtained through interviews with patients and their relatives. Seizure focus lateralization was determined through a comprehensive evaluation of medical history, neurological examination, seizure semiology, video-EEG, and clinical neuroimaging. Detailed clinical data, including location and Engel outcome, is available in ***Supplementary Table S1*.** Twenty-one patients were diagnosed with left-sided TLE. The mean age of seizure onset was 20.42±10.22 years (range=2-49 years), with a mean epilepsy duration of 14.20±10.97 years (range=1-45 years). Eight patients (26%) had a history of childhood febrile convulsion. Based on quantitative hippocampal MRI volumetry ^92^, 20/31 patients (65%%) showed marked hippocampal atrophy ipsilateral to the focus (*i.e.*, absolute ipsilateral-contralateral asymmetry z-score > 1.5 and/or ipsilateral volume z-score < -1.5). At the time of the study, 14/31 patients underwent resective temporal lobe surgeries. In those, post-surgical seizure outcome was assessed using Engel’s modified classification^93^, with an average follow-up duration of 23±15.50 months. Post-surgery, 9 patients (64%) achieved complete seizure freedom (Engel-I), and 5 patients (36%) had recurrent seizures. A total of 9 specimens were available for histopathological analysis. Here, 4 patients showed mesiotemporal/hippocampal sclerosis, one patient had evidence of mesiotemporal ganglioglioma, and 4 for mild dysplasia in the mesiotemporal structures. There were no significant differences in males/females (χ^2^=0.20, p=0.66) and there were no significant differences in age (*t*=-1.01, *p*=0.32) between patients and controls. The study was conducted in accordance with the Declaration of Helsinki, and the MRI data acquisition protocols were approved by the Research Ethics Board of the Montreal Neurological Institute and Hospital. All participants provided informed consent.

### Declarative memory paradigm

All participants underwent episodic and semantic retrieval fMRI tasks (**Figure 1**). These tasks were carefully designed to be closely matched *i.e.,* they were based on (1) equivalent symbolic stimuli to accommodate a bilingual participant pool in the Montreal area; (2) the same number of trials that were presented in a pseudo-randomized manner; (3) an identical task structure with three alternative forced choice response at the retrieval phases; and (4) two difficulty levels.

A) **Episodic Memory.** The task had two phases. During the encoding phase, participants were shown paired images of objects and asked to memorize them. To ensure that the images were well-matched, we used the UMBC similarity index^94^ to control for semantic relatedness. Specifically, we selected items where the similarity score remained below 0.3 on a scale of 0 to 1. The task comprised 56 pseudo-randomized trials, with 28 difficult (with paired images shown only once) and 28 easy (paired images shown twice) trials. In the retrieval phase, which took place after several minutes of delay, participants were presented with an image of an object at the top of the screen (*prime*) and three different objects at the bottom. From these choices, subjects were asked to identify the *target* that was paired with the *prime* during encoding. We only analyzed data from the retrieval phase.
B) **Semantic Memory.** The semantic task involved a retrieval phase only. In each trial, participants were presented with an image of an object at the top of the screen (*prime*) and three different objects at the bottom (1 *target*, 2 *foils*). From these choices, subjects were asked to identify the item (*target*) that was most conceptually related to the *prime*. The 56 pseudo-randomized trials were also modulated for difficulty (*i.e.,* there were 28 difficult trials where prime/target shared a similarity index ≥0.7, whereas prime/foils shared a similarity between 0.3 and 0.7, as well as 28 easy trials, where prime/target shared a similarity ≥0.7 and prime/foils shared a similarity between 0 and 0.3).
C) **Performance metrics**. Individual episodic and semantic performance was computed by averaging the scores in the difficult and easy trials. We also averaged reaction time.

**Figure 1.**
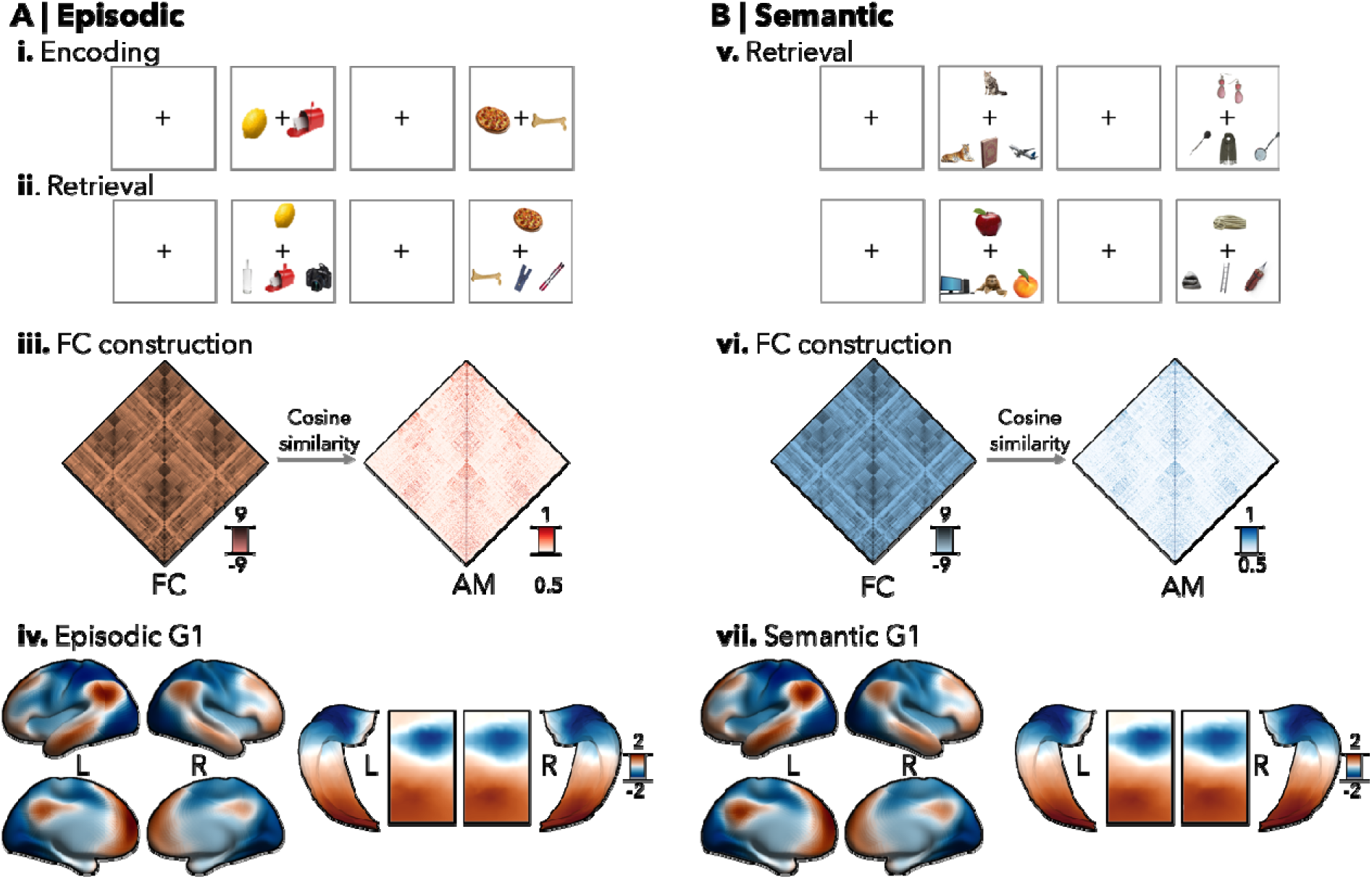
Episodic and Semantic memory tasks. **A**. The episodic memory task included encoding **(i)** and retrieval phases **(ii)**, while **(B) the** semantic memory task only included a retrieval phase **(v)**. Both tasks were matched in terms of stimuli, number of trials, difficulty levels, and task structure, such that retrieval phases included three alternative forced-choice responses. Functional connectomes (FC; **iii, vi**) were independently derived from episodic and semantic task FC and affinity matrices (AM) were built based on cosine similarity, followed by gradient decomposition. (**iv, vii).** Neocortical and hippocampal principal gradients (G1) projected to the corresponding surfaces.

### Additional screening for executive function and general cognitive impairment

Participants also completed the MoCA^95^ tests outside the scanner. The paradigm has been developed to detect mild cognitive impairment and dementia.

### MRI Acquisition

MRI data were acquired on a 3T Siemens Magnetom Prisma-Fit with the 64-channel head coil. T1-weighted scans were acquired with a 3D-magnetization-prepared rapid gradient-echo sequence (MPRAGE; 0.8 mm isovoxels, matrix=320×320, 224, sagittal slices, TR=2300 ms, TE=3.14 ms, TI=900 ms, flip=9°, iPAT=2, partial Fourier=6/8, FOV=256×256 mm^2^). The DWI data was obtained with a 2D spin-echo echo-planar imaging sequence with multi-band acceleration and consists of three distinct shells, each with different b-values 300, 700, and 2000s/mm^2^, with each shell acquired with 10, 40, and 90 diffusion weighting directions, respectively (1.6 mm isotropic voxels, TR=3500LJms, TELJ=64.40 ms, flip angleLJ=LJ90°, refocusing flip angle=180°, FOV=224×224 mm^2^, slice thickness=1.6 mm, multi-band factor=3, echo spacing=0.76 ms). All fMRI were acquired with a 2D/BOLD echo-planar imaging sequence (3.0 mm isovoxels, matrix=LJ80×80, 48 slices oriented to AC-PC-30 degrees, TR=600 ms, TE=30 ms, flip=50°, FOV=240×240 mm^2^, slice thickness=3 mm, multi-band factor=6, echo spacing=0.54 ms). During the resting-state functional MRI (rs-fMRI), participants were instructed to fixate on a grey cross and not think of anything. All fMRI scans were presented via a back-projection system and responses were recorded using an MRI-compatible four-button box. The episodic and semantic retrievals both lasted approximately 6 minutes.

### MRI Processing

Scans were pre-processed using *micapipe*^96^ *version 0.2.2* (https://github.com/MICA-MNI/micapipe), an open-access multimodal preprocessing, surface-mapping, and data fusion software. T1-weighted data were de-obliqued and reoriented to ensure a consistent orientation across images. Next, a linear coregistration, intensity non-uniformity correction, and skull stripping were applied to extract the brain region from the surrounding non-brain tissues. DWI data underwent denoising, eddy current-induced distortion correction, and motion correction as well as non-uniformity bias field adjustment. The corrected DWI data was fitted with a diffusion tensor model,^97^ allowing for the computation of mean diffusivity images (MD).^98^ Cortical surfaces were extracted with FreeSurfer *6.0* from each T1w scan, followed by manual surface generation. Resting-state and task-fMRI were preprocessed using a combination of FSL^99^ *6.0*, ANTs^100^ *2.3.4*, and AFNI^101^ *20.3.0* software and involved the following steps: *(1)* removal of first five TRs, *(2)* image reorientation, *(3)* motion correction, *(4)* distortion correction based on AP-PA blip field maps, *(5)* high-pass filtering to be above 0.01Hz, *(6)* MELODIC decomposition, *(7)* Linear/Non-linear co-registration to the T1-weighted scans, *(8)* nuisance signal regression with FMRIB’s ICA-based Xnoiseifier^102^ (ICA-FIX), *(9)* native cortical surface registration, *(10)* surface-based registration to the Conte69 surface template with 32k surface points (henceforth, *vertices*) per hemisphere, *(11)* surface-based smoothing with a Gaussian diffusion kernel with a full-width-at-half-maximum, FWHM, of 10 mm, and *(12)* statistical regression of motion spikes. The hippocampus and surrounding structures were automatically segmented using HippUnfold^92^ *version 1.0.0,* (https://hippunfold.readthedocs.io/). The task-fMRI and MD data were then sampled along the hippocampal midthickness surface and FWHM=2 mm surface smoothing was applied.

### Gradient identification and alignment

For computational efficiency, surface-mapped neocortical time series were downsampled to a mesh with 10k vertices using *connectome workbench*.^103^ Intrahemispheric neocortical functional connectomes were generated for each functional state (episodic, semantic, rest) by calculating Pearson correlations between the time series for all vertices for each participant, resulting in a 5k-by-5k connectome per hemisphere (**Figure 1**). As for the neocortex, internal hippocampal connectomes were generated from the extracted timeseries resulting in a 419 by 419 functional connectivity matrix per hippocampus. We applied Fisher r-to-z transformations to both neocortical and hippocampal functional connectomes and retained only the top 10% of row-wise connections. We constructed affinity matrices using a cosine similarity kernel to measure the similarity of connectivity patterns between regions. Diffusion map embedding,^65,104^ a non-linear dimensionality reduction technique implemented in *Brainspace*^105^ *version 0.1.3* (https://brainspace.readthedocs.io/; **Figure 1**), was applied to identify low-dimensional eigenvectors (*i.e.,* gradients) that accounted for the variance in neocortical and hippocampal connectomes. Diffusion mapping is a powerful and efficient tool to compress large-scale datasets. This algorithm uses the diffusion operator *P*α to create a new representation of data and is guided by two key parameters: α, which controls the influence of sampling point density on the manifold, and, *t*, which represents the scale of the data. To maintain global connections among data points in the embedded space, α was set to 0.5 (α*=0, maximal influenc*e, α*=1 no influence)* and *t* was set at 0, consistent with previous studies.^65,105^ To ensure that results were consistent with the prior literature that derived functional gradients mainly from rs-fMRI data,^65,70^ we first generated intra-hemispheric reference gradients by averaging rs-fMRI connectomes across groups and aligning them to a normative gradient template derived from the rs-fMRI data of the Human Connectome Project.^106^ We used Procrustes rotation to align individual and state-specific task-fMRI gradients with these normative reference gradients, thereby ensuring that gradients were consistent in ordering and sign across participants and memory states^107^. Similar procedures were performed to align hippocampal gradients. Finally, we normalized neocortical and hippocampal gradient scores relative to the distribution in controls, and sorted gradients in patients relative to the hemisphere ipsilateral to the seizure focus, as in prior work.^87^

### Statistical Analysis

After gradient alignment, we inferred between-group differences using surface-based linear models implemented in *Brainstat*^108^ *version 0.4.2* (https://brainstat.readthedocs.io). We additionally controlled for age and sex and corrected for vertex-wise multiple comparisons using random field theory for non-isotropic imaging,^109^ at a set family-wise error of p_FWE_<0.05. We computed the association between gradient scores in regions showing significant between-group differences and cognitive scores. To investigate the association between gradients and memory, we examined the gradient scores while controlling for MoCA which indicated mild cognitive impairments. Finally, statistical mediation analyses tested whether the association between group and episodic memory performance is transmitted via gradients.

### Morphological and microstructural substrate analysis

To assess structural substrates of functional network changes, we also compared neocortical and hippocampal thickness and mean diffusivity (MD) between controls and TLE patients. These features have previously been utilised to demonstrate both hippocampal as well as neocortical alterations in TLE. ^35,110,111^ Neocortical thickness was quantified as the Euclidian distance between corresponding pial and white matter vertices, and we used mid-thickness surfaces to sample MD from co-registered diffusion MRI data. To establish an accurate spatial correspondence, cortical thickness and MD maps were registered to the Conte69 surface template and were subsequently downsampled from 32k vertices to 5k vertices/hemisphere. Hippocampal thickness and hippocampal MD was measured using a similar method, using hippocampal surface meshes derived with HippUnfold^92^ *v1.0.3,* (https://hippunfold.readthedocs.io/). Multivariate surface-based linear models then assessed morphological and microstructural differences between patients and controls in both hippocampal and neocortical regions, while controlling for effects of age and sex. Finally, we investigated the association between atypical morphology patterns and microstructure and observed TLE-related gradient alterations. Specifically, we computed the mean thickness and MD scores in regions showing significant functional G1 differences between controls and TLE patients, and correlated these measures with G1 changes.

### Functional decoding via task-based meta-analysis

We also conducted a functional decoding analysis based on Neurosynth ^26^ (https://neurosynth.org/), a platform that allows for large-scale *ad hoc* meta-analysis of task-based fMRI data. Here we studied the spatial associations between TLE-related functional gradient changes and previously published fMRI activations to identify cognitive terms that elicit similar activation patterns.

### Data availability

The Human Connectome Project dataset used for generating normative resting-state gradients is available at https://db.humanconnectome.org/. Our healthy control cohort is made up of a subset of participants from the MICA-MICs^112^ dataset which is openly available on the Canadian Open Neuroscience Platform data portal (https://portal.conp.ca/dataset?id=projects/mica-mics) and Open Science Framework (https://osf.io/j532r/). Functional connectome gradients will be available on <u>osf.io</u>.

## Results

Hippocampal and neocortical surface models for each participant were generated using automatic segmentation^92,96^ procedures based on T1-weighted MRI. Functional data were co-registered to these surfaces, and resulting neocortical and hippocampal timeseries were used to derive functional connectomes for resting, episodic, and semantic memory states for each participant. Diffusion map embedding,^105^ a non-linear dimensionality reduction technique, was applied to estimate low-dimensional eigenvectors explaining spatial gradients of connectivity variance. For consistency, reference gradients were derived from rs-fMRI data^106^ and subject- and state-specific neocortical and hippocampal gradients were aligned to these using Procrustes rotations, as in previous work ^105,107^ (for pre-alignment gradient patterns, please see ***Supplementary Figure S1***).

### Declarative memory topographies

Studying neocortical connectivity in healthy individuals, we derived the first eigenvectors (G1) during both episodic and semantic states separately (for findings of subsequent gradients, please see ***Supplementary Figure S2***). In episodic/semantic states, G1 explained 22/21% of functional connectome variance and described a sensory-to-transmodal neocortical gradient, as expected from previous work in healthy adults based on rs-fMRI connectivity.^65^ Studying hippocampal networks in controls, we also derived the first eigenvectors in episodic and semantic memory states (**Figure 1**). In episodic/semantic states, G1 accounted for 40/38% of functional connectome variance and described a canonical postero-anterior spatial pattern.^76^ State-specific neocortical and hippocampal gradients spatially correlated in healthy controls (*r*=0.99, *p*=0, *n_perm_*=1000), suggesting that both memory systems were captured by both sensory-transmodal neocortical and posterior-anterior MTL trends.

### Atypical task-based gradients in patients with MTL pathology

Comparing neocortical topographies in TLE patients to controls, we found reduced G1 scores in the episodic memory state in patients (*p_FWE_*<0.05, mean effect sizes in significant regions *d*>0.50; **Figure 2A**). Functional G1 alterations were primarily localized in the ipsilateral lateral temporal and bilateral posterior cingulate regions. Between-group differences were also seen in the semantic memory state (**Figure 2B**), localized in the bilateral lateral temporal and ipsilateral angular gyrus. The TLE-related functional G1 contractions are indicative of a functional dedifferentiation between these unimodal and transmodal systems. Consistent with this finding, the histogram depicting overall mean gradient scores, showed an overall reduction in the spread of gradient scores in TLE patients relative to controls (**Figure 2**). Considering the hippocampus, conversely, we observed an expansion of G1 scores in the episodic memory state when comparing patients to controls, which was particularly significant in anterior divisions ipsilaterally (*p_FWE_*<0.05, *d*=-0.84). The expanded functional G1 in the anterior portions reflects increased connectivity profile variability in TLE, indicating a great intra-hippocampal functional disconnection. In other words, the anterior hippocampal gradient scores in TLE are shifted away from the gradient midpoint, further increasing inter-regional differentiation with the posterior hippocampus. While semantic G1 expansion was also observed in TLE, no vertices survived multiple comparison correction.

**Figure 2.**
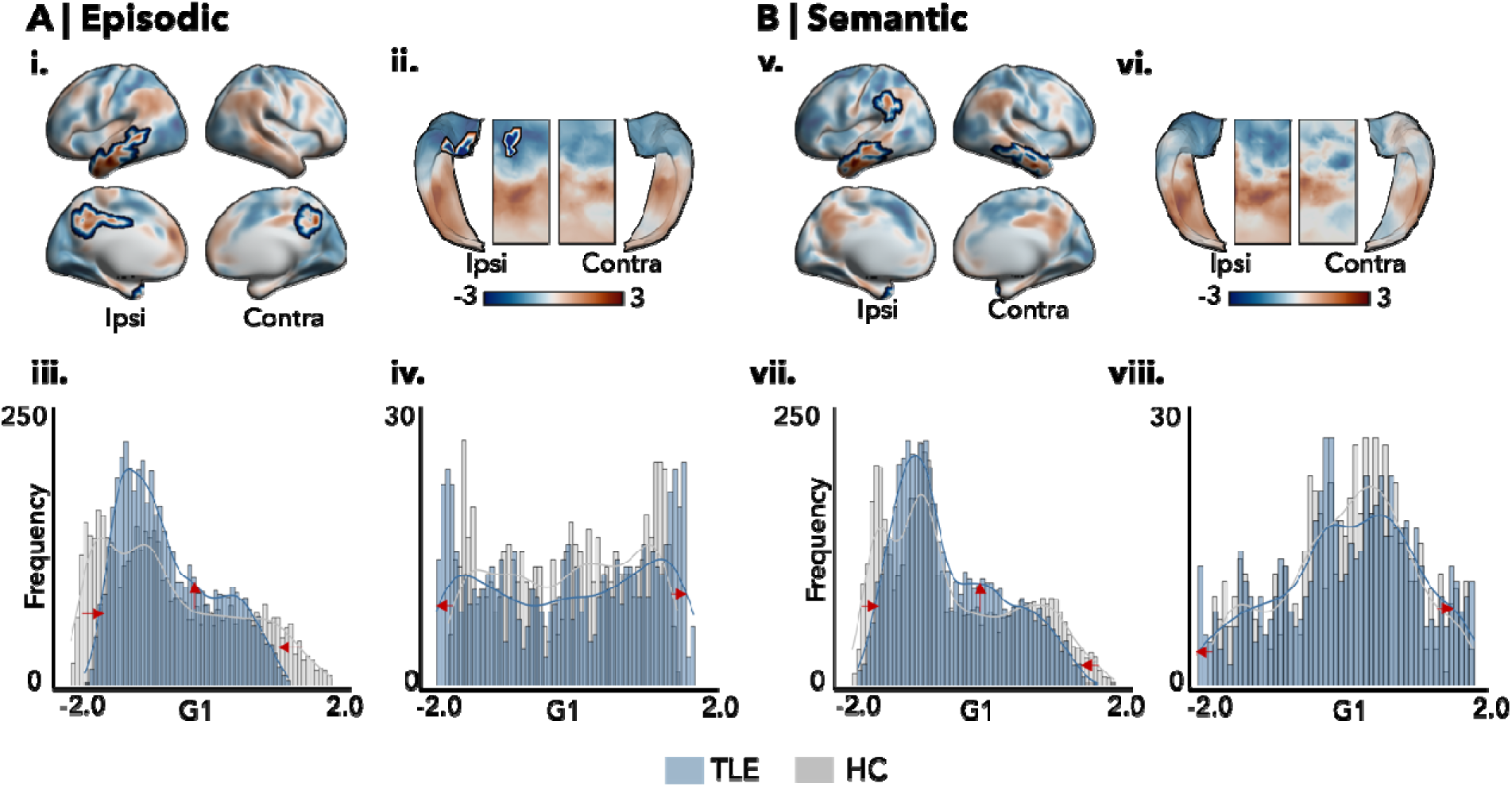
Gradient alterations in patients with MTL pathology. Group differences in neocortical **(i,v)** and hippocampal **(ii, vi)** gradient scores for episodic **(A)** and semantic states **(B).** Significant *p*-values were highlighted after correcting for multiple comparisons (*p_FWE_*<0.05, black outlines). Histogram plot of neocortical gradient contractions **(iii,vii)** and hippocampal gradient expansions **(iv,viii)** in TLE patients relative to controls in episodic and semantic states, respectively. **Abbreviations:** HC=healthy controls, TLE= temporal lobe epilepsy, G1 = Gradient 1, G2= Gradient 2.

### Relation to temporo-limbic structural alterations

An additional analysis explored how proxies of TLE-related structural pathology^35^ contribute to episodic functional memory network reorganization. Proxies were derived from neocortical and hippocampal MRI measures of grey matter (*i.e.,* cortical thickness) and diffusion parameters (*i.e.,* cortical mean diffusivity, MD). Comparing patients and controls using multivariate models based on these parameters, significant alterations were observed in patients (*p*<0.05, **Figure 3A, B;** for univariate findings, see ***Supplementary Figure S3***), mainly localized in ipsilateral tempo-limbic areas and anterior regions of the ipsilateral hippocampus. We next examined whether these structural disruptions are associated with previously observed functional alterations (**Figure 3C**). To this end, we computed the overall temporo-limbic structural changes in TLE patients and controls, both at the level of the neocortex and hippocampus and conducted a correlation analysis. This analysis revealed an association between overall episodic functional G1 changes and structural alterations (*r*=-0.22, *p_FDR_*=0.03). Further tests showed a slightly more selective association with hippocampal (r=-0.22, *p_FDR_*=0.04) than with neocortical functional G1 measures (*r*=0.19, *p_FDR_*=0.08).

**Figure 3.**
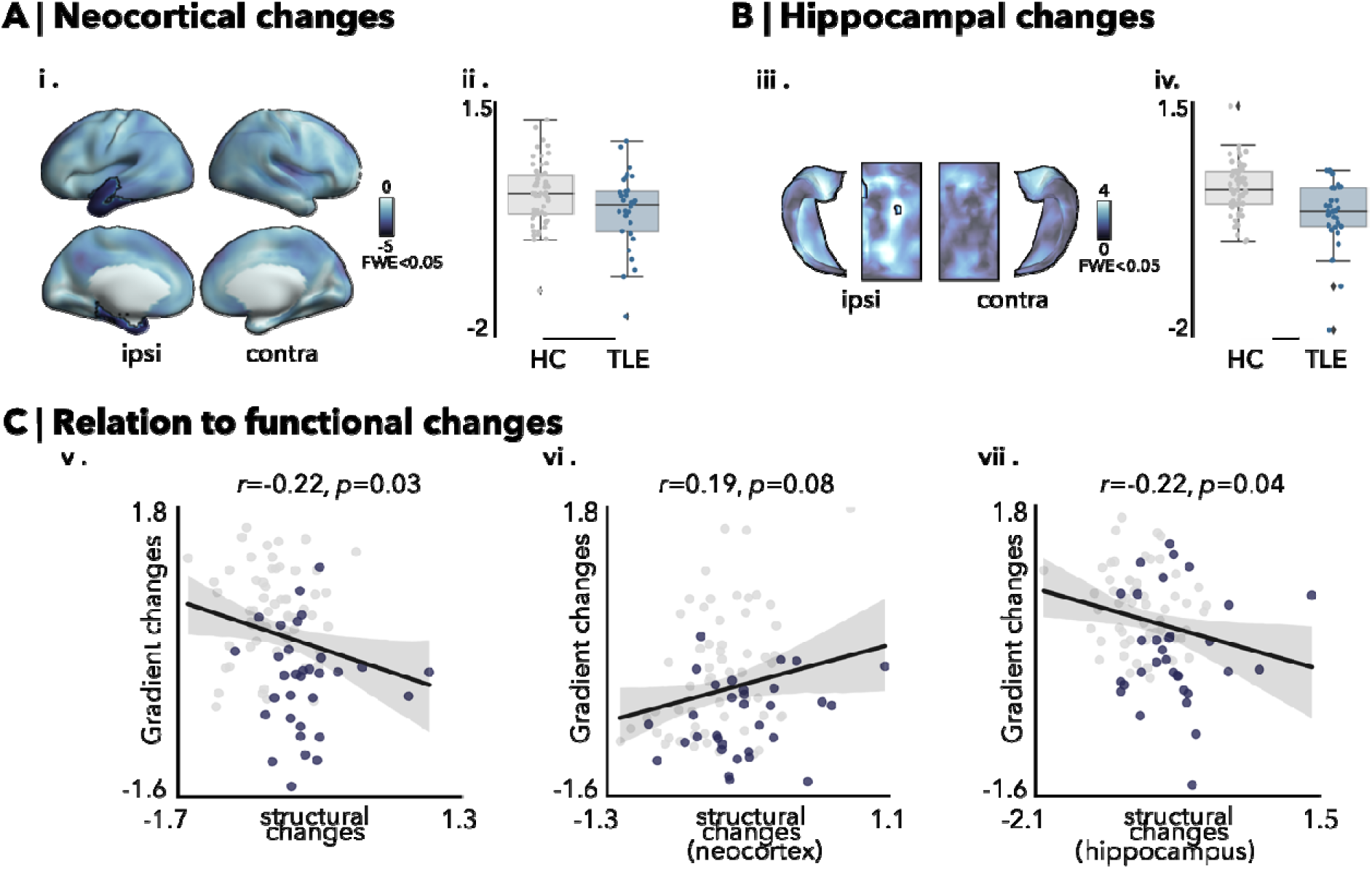
Associations to temporo-limbic pathology. **A-B)** Differences in neocortical **(i)** and hippocampal **(iii)** MRI-proxies of pathology (using a multivariate aggregate of cortical thickness and mean diffusivity) between TLE patients and controls. Regions showing alterations in TLE are highlighted (p_FWE_<0.05, black outline), with mean values represented in **(ii, iv). C**) FDR-corrected relationship between overall episodic changes to **(v)** overall structural changes, **(vi)** structural changes in the neocortex, and, **(vii)** hippocampus (HC=grey; TLE=dark purple). **Abbreviations:** ipsi= ipsilateral, contra=contralateral, HC=healthy controls, TLE=temporal lobe epilepsy.

### Relation to behavioural indices of declarative memory

We examined the relations between participants’ behavioural performance and functional memory network reorganization. Behaviorally, patients showed markedly reduced episodic memory accuracy relative to controls (*t*=-4.40, *p*<0.001, *d*=-0.97), and only subthreshold impairment in semantic memory (*t*=-1.42, *p*=0.16, *d*=-0.31). We similarly observed faster reaction times for accurate recall in controls relative to patients (***Supplementary Figure S5***).

Notably, functional G1 scores in both neocortical (*r_ipsi_*=0.29, *p_FDR_*=0.02; r_contra_=0.27, *p_FDR_*=0.02) and hippocampal (*r_ipsi_*=0.29, *p_FDR_*=0.02) clusters of significant between-group differences (see *Figure 2*) correlated with episodic memory performance (**Figure 4A**). *Ad hoc* meta-analytical decoding^26^ confirmed that TLE-related gradient contractions in the episodic state were enriched for memory-related terms, supporting the behavioural associations (***Supplementary Figure S6***). In semantic memory states, hippocampal functional; G1 (*r_ipsi_*=0.21, *p_FDR_*=0.06) correlated with semantic task performance at trend levels, but not neocortical functional G1 (*r_ipsi_*=0.02, *p_FDR_* =0.85; r_contra_=-0.03, *p_FDR_*=0.95; **Figure 4B**).

**Figure 4.**
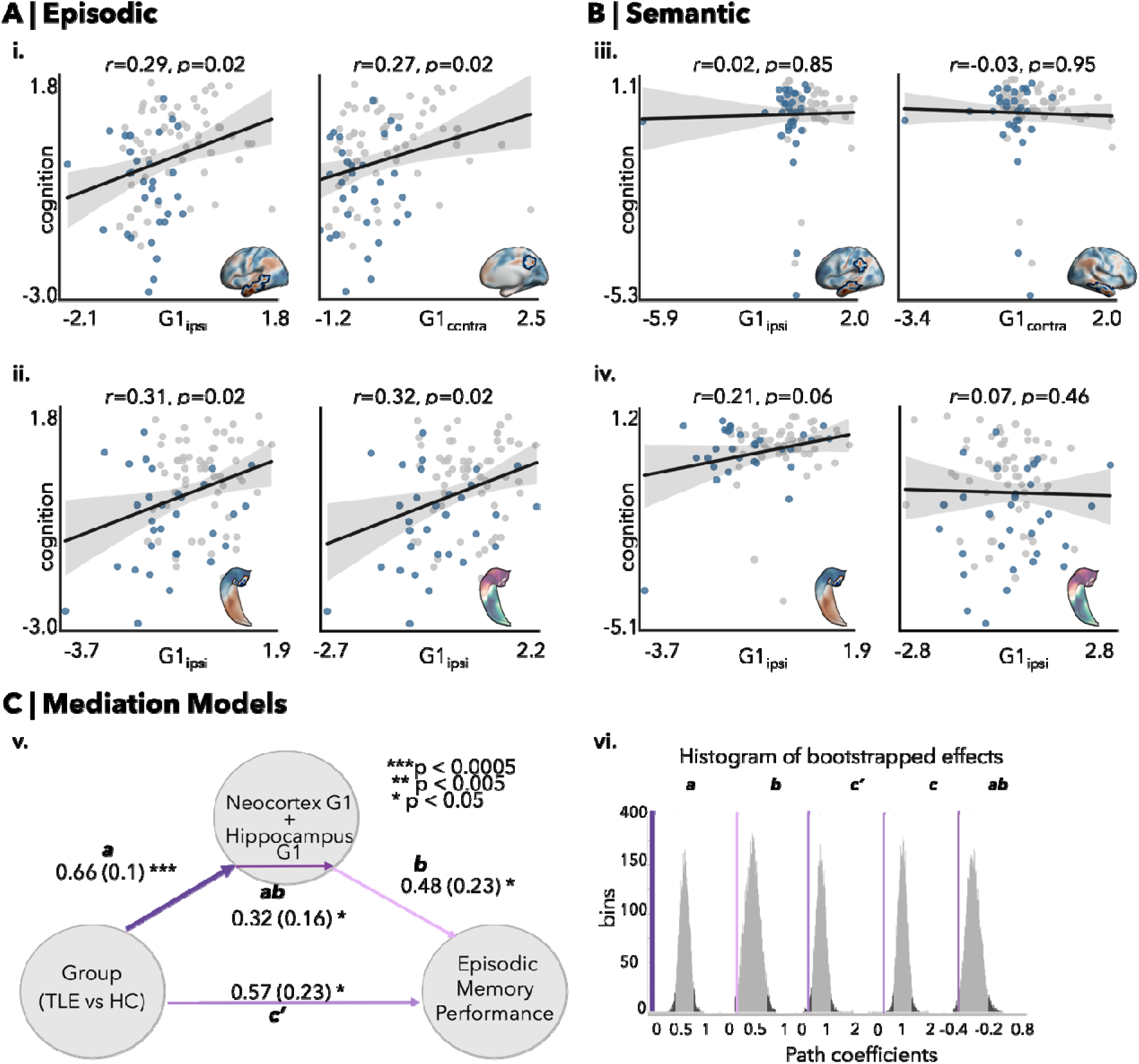
Behavioral associations. Mean G1 scores of regions showing significant case-control differences in the ipsilateral and contralateral hemispheres for neocortical and hippocampal G1 in episodic **(A)** and semantic **(B)** memory states, correlated with overall behavioural performance. Controlling for the aggregate effects of morphology and superficial white matter properties (mean diffusivity), hippocampal episodic G1 (**ii;** *right panel)* correlated significantly with task performance, while hippocampal semantic G1 (**iv;** *right panel)* correlation to memory performance disappeared. (HC=grey; TLE=blue). **C.** Statistical mediation analysis revealed the relationship between the group and episodic memory performance, transmitted via overall neocortical G1+ hippocampal G1_ipsi_ **(v)** scores. **(vi)** Histogram of effects (nboot=1000). **Abbreviations:** G1 = Gradient 1, Ipsi= ipsilateral, Contra= contralateral, a = Group ∼ Gradient scores, b= Gradient scores ∼ Episodic memory performance, c’= Group ∼ Episodic memory performance(residual), ab= mediation effect.

To assess specificity, we also administered the MoCA test to probe mild cognitive dysfunction. Patients with TLE showed reduced MoCA (*t*=-3.37, *p*=0.001, *d*=-0.74) scores relative to controls. Controlling for individual MoCA scores, significant functional G1 changes persisted in our cohort, in both the hippocampus (*p_FWE_*<0.05, *d*=-0.80, 4% increase in mean effect size) and neocortex (*p_FWE_*<0.05, *d*=-0.50).

In a final analysis, we assessed whether functional G1 alterations mediated the relationship between group (TLE *vs* controls) and episodic memory performance (**Figure 4C**, for individual mediation, see ***Supplementary Table S2***). To this end, we first computed overall episodic alterations by averaging G1 scores in significant regions and submitted this to a statistical mediation analysis. Group and functional G1 alterations correlated strongly (a, p<0.0005), as did functional G1 alterations and episodic memory performance (b, p<0.05), and group and episodic memory performance (c’, p<0.05). Importantly, functional G1 alterations in hippocampal and neocortical regions mediated the relationship between group and episodic memory performance (ab, p<0.05). Overall, our statistical mediation analysis, thus, revealed that the relationship between group and episodic memory performance is transmitted via neocortical and hippocampal functional gradients.

## Discussion

This study demonstrated *(i)* a selective reorganization of both neocortical and hippocampal episodic functional memory networks in patients with TLE, *(ii)* a close association between functional reorganization and hippocampal, but not neocortical *in vivo* proxies of MTL pathology, and *(iii)* a contribution of atypical temporo-limbic functional organization to behavioural deficits in TLE patients. Our study harnessed several innovative elements, notably closely matched functional MRI tasks of episodic and semantic memory administered in TLE patients and controls, state-of-the-art MRI segmentation and multimodal image preprocessing methods, as well as connectivity gradient mapping techniques to identify functional topographies in a data-driven manner. These techniques were applied to a cohort of TLE patients with variable degrees of MTL pathology and memory impairment, allowing for the study of structure-function relationships in declarative memory systems in the human brain. By identifying the functional topographic alterations of episodic memory networks in TLE that mediate cognitive impairment and their structural underpinnings, our work emphasizes a key role of the MTL in episodic over semantic memory processes.

The segregation of episodic and semantic memory systems^3^ supported by our findings is in line with foundational neuropsychological investigations dissociating list recall performance for episodic memory and picture naming for semantic memory.^38,60,61^ Moreover, task-based neuroimaging studies have pointed to distinct neural substrates of these task demands.^9,16^ The memory tasks^46^ we employed were carefully designed to distinctly tap into episodic and semantic memory while being homogenous in low-level task structure. A recent behavioral study from our group supported the utility of these tasks to assess cognition in both healthy and diseased cohorts, showing reduced episodic, but relatively preserved semantic recall in TLE.^46^ Despite these potential differential impairments of TLE patients, mounting evidence suggests that episodic and semantic memory systems may nevertheless share overlapping neural correlates.^9,22,23^ By capturing both distinctive features and potential overlaps in the neural underpinnings of episodic *vs* semantic memory, we employed a data driven approach that mapped functional topographic patterns associated with each of these processes. In particular, we analyzed task-based fMRI data across both episodic and semantic memory states, and mapped resulting task-based connectomes into topographic connectivity gradients.^65,70^ Gradient mapping techniques have previously been applied to several modalities, in particular resting-state fMRI connectivity as well as structural and microstructural MRI measures.^65,70,72,82,113,114^ Instead of performing a conventional functional gradient mapping across task-free fMRI sessions, the current study derived topographic gradients during episodic and semantic memory states separately, and found that they largely followed similar intrinsic organizational axes irrespective of the choice of gradient alignment.^10,23,25^ Specifically, gradients associated to both memory states followed a sensory-transmodal neocortical and posterior-anterior hippocampal pattern, in line with previous observations.^65,70,71,74–76^ Transmodal core regions (*i.e.,* DMN) are recognized to occupy a cortical territory that is maximally distant and separate from unimodal sensory systems, balancing segregated processing streams and integration.^66,68,65,66,115^ This is suggested to help decouple cognition, for instance through self-generated thought processes and episodic representations, from salient sensory information of the immediate external environment.^116–121^ Conversely, in the hippocampus, anterior segments are thought to have broader tuning properties and are suited to generalization, while posterior segments are more narrowly tuned and appear more suited to memorizing specific instances for contextualization.^122,123^ This can be supported by the preferential coupling of anterior divisions of the MTL with transmodal systems such as the DMN, while posterior segments become increasingly connected to unimodal and sensory-motor systems.^75,84,124,125^ Together, our findings echo this unified account of large-scale hierarchical organization of the brain and bridges the dichotomy between different declarative memory systems with reliable correlates to behavioural phenotypes.^77,126,127^

In TLE, previous work has mainly reported impaired memory performance and atypical functional activation patterns in the MTL and beyond.^128,129^ Here, we extend this literature by showing a marked reorganization of task-based functional network topographies during episodic and semantic retrieval states. These effects were strongest in bilateral neocortical regions for both memory states, with selective involvement of the MTL region ipsilateral to the seizure focus on episodic states. Consistent with previous work in TLE based on rs-fMRI data,^128^ we found neocortical gradient compressions in regions susceptible to TLE-related pathology.^128,130^ Other rs-fMRI studies reported simultaneous increases in short- and decreases in long-range connections in temporo-limbic and dorsomedial regions in TLE^128,129^. Moreover, a large body of structural covariance and diffusion MRI studies has shown reduced connectivity of primarily the temporo-limbic diseased epicentre, but also broader brain networks.^35,128^ Studying the hippocampus using gradient mapping techniques, we observed an extended episodic, but not semantic principal gradient in TLE patients relative to controls. These hippocampal gradient alterations are, thus, in an opposing direction than the neocortical findings in TLE. An expanded hippocampal gradient in TLE patients is readily interpretable as an increased functional differentiation between anterior and posterior segments, which could reflect reductions in intra-hippocampal crosstalk along the long axis. In healthy individuals, long axis organization of the hippocampus has previously been suggested to relate to hierarchy of gist *vs* detailed oriented recollection.^127,131, 132^ Studies show that the anterior hippocampus acts as a hub, recruiting transmodal neocortical regions for schema construction, evidenced by direct and reciprocal connections between anterior temporal and dorsal and ventromedial prefrontal cortices.^2,133,134^ Furthermore, the coactivation of posterior hippocampal and unimodal sensory regions of the neocortex is thought to allow detailed elaborations. ^134^ In TLE patients, previous work indicate that patients exhibit a reduced capacity for recalling internal details of specific personal episodes but with relatively preserved narrative components,^135^ suggesting that hippocampus is essential for recollecting sensory perceptual aspects of previous experiences. As such, the increasing spatial disconnection between anterior and posterior hippocampus observed in our TLE patients could reflect a compensatory retrieval process emphasizing key elements over extraneous details during the episodic memory task^135^.

Together, our findings showing the co-occurrence of gradient contractions in neocortical systems with simultaneous gradient expansion of hippocampal subregions may represent a topographic mechanisms of memory reorganization in TLE, compatible with atypical functional segregation in mesiotemporal and neocortical systems, respectively. ^86,128^ Accordingly, neocortical gradient contractions indicate that transmodal regions are relatively dedifferentiated from unimodal systems, likely stemming from imbalances in both local and distant connectivity. On the other hand, expansions in hippocampal connectivity gradients are likely indicative of reductions in intra-hippocampal connectivity, which would contribute to marked divergences in signalling between anterior and posterior divisions. As such, our findings echo task-fMRI studies showing widespread extratemporal connectivity reductions and intrinsic MTL connectivity increases,^47^ and less concordant fMRI activations between hippocampal long axis and sensory-transmodal gradients in TLE relative to controls.^32^ Collectively, the neocortical gradient contraction and hippocampal gradient expansion reported here provide a compact signature of reorganization of declarative memory systems in TLE.

Prior studies have reported atypical microstructure and morphology in TLE in both the MTL as well as adjacent temporo-limbic neocortical systems, ^47, 34,35,89,90,136–138^ and have suggested that such changes could alter the spatial configuration of functional networks and influence hierarchical organization supporting episodic memory function^87^. In line with previous work,^35,111,128^ white matter disruptions and atrophy were observed in our TLE patients, with most marked findings in temporo-limbic regions and hippocampal anterolateral regions ipsilateral to the seizure focus. These findings could reinforce earlier models that propose increased susceptibility of paralimbic regions to structural and functional changes, potentially due to a greater capacity for plasticity and connectivity rearrangement.^88,139^ Interestingly, while neocortical structural changes did not reflect functional changes, morphological and microstructural derangements of the hippocampus correlated with episodic memory network reorganization. Previous studies have consistently revealed anterior hippocampal atrophy^140,141^ and alterations in tissue microstructure^128^ in TLE patients, aligning marked neuronal loss in the anterior hippocampus.^142^ Such structural damage in crucial hubs like the hippocampus extend beyond localized effects and can compromise communication within the interconnected memory network^140,143^. The functional association with pathological markers selective to the hippocampus reported herein, therefore, provides specificity, and reinforces the key role of hippocampal integrity in episodic memory network organization.^144,145^

Previous studies have also linked network alterations in TLE patients to cognitive impairment.^47,87,129,136,138^ Consistent with a recent study from our group ^46^, episodic task impairments were marked in TLE, while semantic task performance remained intact. Additionally, neocortical, and hippocampal gradient alterations were found to relate to episodic impairments in TLE. While we acknowledge that our conservative patient inclusion criteria resulted in a relatively modest sample size of 31 TLE patients, the observed memory deficits and functional perturbations were unlikely influenced by neurodegenerative disorders,^146,46^ as functional alterations persisted even after controlling for overall measures of cognitive function (*i.e.* MoCA). Behavioural indices of cognitive impairment in TLE were mediated by functional changes, confirming the already well-established literature on episodic impairment in TLE. In line with the hierarchical models of memory,^147,148^ our findings lend support to the idea that semantic memory relies more on extra-hippocampal brain regions compared to hippocampal-dependent episodic systems. This is consistent with evidence from TLE patients who underwent anterior lobe resections,^59^ and corroborates the controlled semantic cognition framework with the ATL acting as a central semantic hub.^1,16,29^ Additionally, our findings are accounted for by the complementary learning systems framework,^149,150^ which stipulates that hippocampus-encoded semantic memories undergo a long-term consolidation process^151,152^ that renders the memory trace independent from the hippocampus, and thus more resilient to its structural pathology. Likewise, the multiple trace theory^57,58^ specifies that episodic memories are consistently mediated by the hippocampus allowing a vivid recollection of context-rich events,^58,153^ and each instance of recalling an episodic memory triggers subsequent re-encoding, leading to the formation of multiple traces or engrams, that are mediated by hippocampo-neocortical neuron ensembles.^57,58,153,154,155,156^ In contrast, successful semantic memory recall does not require rich contextual detail and can, therefore, be solely supported by extra-temporal regions.^157^

Collectively, our work presents a novel decomposition analysis of task-fMRI data revealing an atypical and state-dependent reorganization of declarative memory systems in TLE, with a consistent correlation with behavioural impairments and *in vivo* markers of hippocampal pathology. Our findings support a more selective role of the hippocampus in episodic processes, in line with the episodic theory of memory organization ^31,158^. This study also provides insights into mechanisms of cognitive impairments in conditions associated with MTL pathology, showing in particular an opposing pattern of decreased neocortical functional differentiation together with increased intra-hippocampal differentiation. As many drug-resistant patients from this cohort undergo resective MTL surgery, future work could investigate whether gradient patterns will help predict post-operative memory deficits, and thus enrich patient-specific clinical decision-making.

## Acknowledgements

We would like to thank all the TLE patients and controls who participated in this study.

## Supplementary material

Supplementary material is available at *Brain* online.

## Supplementary Materials

**Table S1.** Patient’s demographics and clinical features. **Abbreviations:** L=left; R=right, HD= Handedness

| **ID** | **Age** | **Sex** | **HD** | **Laterality** | **Onset** | **Duration**<br>**(years)** | **Febrile**<br>**convulsions** | **Engel outcome** |
| --- | --- | --- | --- | --- | --- | --- | --- | --- |
| P01 | 54 | M | R | L | temporal | 37 | 0 | n/a |
| P02 | 26 | F | R | L | temporal | 10 | 0 | 1a |
| P03 | 34 | M | R | L | temporal | 4 | 0 | 1a |
| P04 | 20 | F | R | L | temporal | 5 | 0 | 1a |
| P05 | 25 | F | R | L | temporal | 12 | 0 | 1b |
| P06 | 33 | F | R | L | temporal | 6 | 0 | 1a |
| P07 | 57 | M | R | L | temporal | 39 | 0 | 1a |
| P08 | 41 | F | R | R | temporal | 25 | 2 | 1b |
| P09 | 31 | M | R | L | temporal | 14 | 1 | n/a |
| P10 | 52 | M | R | L | temporal | 3 | 0 | n/a |
| P11 | 55 | F | R | L | temporal | 13 | 1 | 1a |
| P12 | 26 | F | L | L | temporal | 9 | 1 | n/a |
| P13 | 25 | M | R | R | temporal | 23 | 0 | n/a |
| P14 | 22 | F | R | R | temporal | 12 | 0 | 2b |
| P15 | 53 | F | R | L | temporal | 45 | 0 | n/a |
| P16 | 24 | M | R | L | temporal | 6 | 0 | n/a |
| P17 | 38 | M | R | L | temporal | 12 | 1 | n/a |
| P18 | 43 | M | R | R | temporal | 13 | 2 | 1a |
| P19 | 32 | M | R | R | temporal | 14 | 0 | 1a |
| P20 | 41 | M | R | L | temporal | 27 | 0 | 2b |
| P21 | 27 | F | R | L | temporal | 12 | 2 | n/a |
| P22 | 25 | F | R | R | temporal | 8 | 0 | 3a |
| P23 | 28 | M | R | L | temporal | 1 | 0 | n/a |
| P24 | 29 | M | R | R | temporal | 7 | 0 | n/a |
| P25 | 27 | F | R | L | temporal | 14 | 0 | n/a |
| P26 | 39 | F | R | R | temporal | 27 | 0 | 1a |
| P27 | 36 | F | R | L | temporal | 9 | 2 | n/a |
| P28 | 18 | F | R | L | temporal | 4 | 0 | n/a |
| P29 | 28 | F | R | L | temporal | 5 | 0 | n/a |
| P30 | 29 | F | R | R | temporal | 11 | 0 | n/a |
| P31 | 56 | M | R | R | temporal | 14 | 0 | n/a |

**Table S2.**
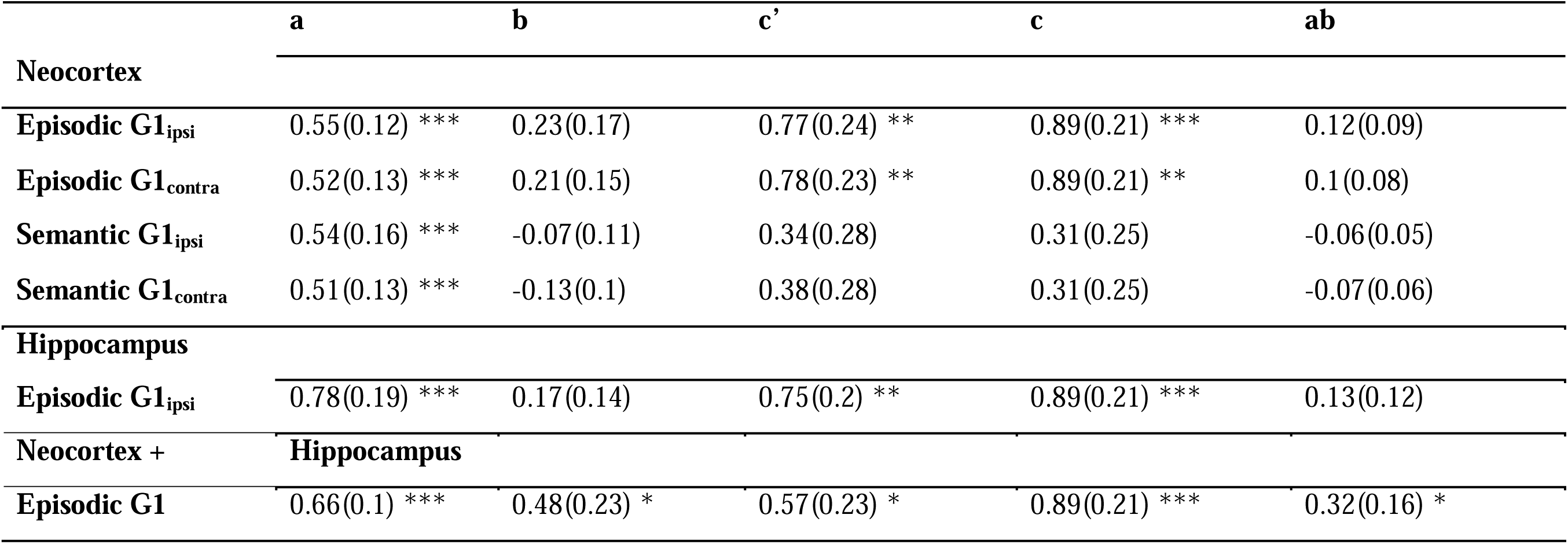
Individual mediation analysis for Episodic and Semantic memory G1. The mean of significant regions showing significant episodic and semantic gradient scores were used as mediators for each analysis. **Abbreviations:** G1=Gradient1, Ipsi=ipsilateral, Contra=contralateral, a = Group ∼ Gradient scores (standard errors in parenthesis), b= Gradient scores ∼ Task performance, c’= Group ∼ Task performance, c’= Group ∼ Task performance (residual), ab= mediation effect

|  | **a** | **b** | **c'** | **c** | **ab** |
| --- | --- | --- | --- | --- | --- |
| **Neocortex** |  |  |  |  |  |
| **Episodic G1<sub>ipsi</sub>** | 0.55(0.12) *** | 0.23(0.17) | 0.77(0.24) ** | 0.89(0.21) *** | 0.12(0.09) |
| **Episodic G1<sub>contra</sub>** | 0.52(0.13) *** | 0.21(0.15) | 0.78(0.23) ** | 0.89(0.21) ** | 0.1(0.08) |
| **Semantic G1<sub>ipsi</sub>** | 0.54(0.16) *** | -0.07(0.11) | 0.34(0.28) | 0.31(0.25) | -0.06(0.05) |
| **Semantic G1<sub>contra</sub>** | 0.51(0.13) *** | -0.13(0.1) | 0.38(0.28) | 0.31(0.25) | -0.07(0.06) |
| **Hippocampus** |  |  |  |  |  |
| **Episodic G1<sub>ipsi</sub>** | 0.78(0.19) *** | 0.17(0.14) | 0.75(0.2) ** | 0.89(0.21) *** | 0.13(0.12) |
| **Neocortex + Hippocampus** |  |  |  |  |  |
| **Episodic G1** | 0.66(0.1) *** | 0.48(0.23) * | 0.57(0.23) * | 0.89(0.21) *** | 0.32(0.16) * |

**Figure S1.**
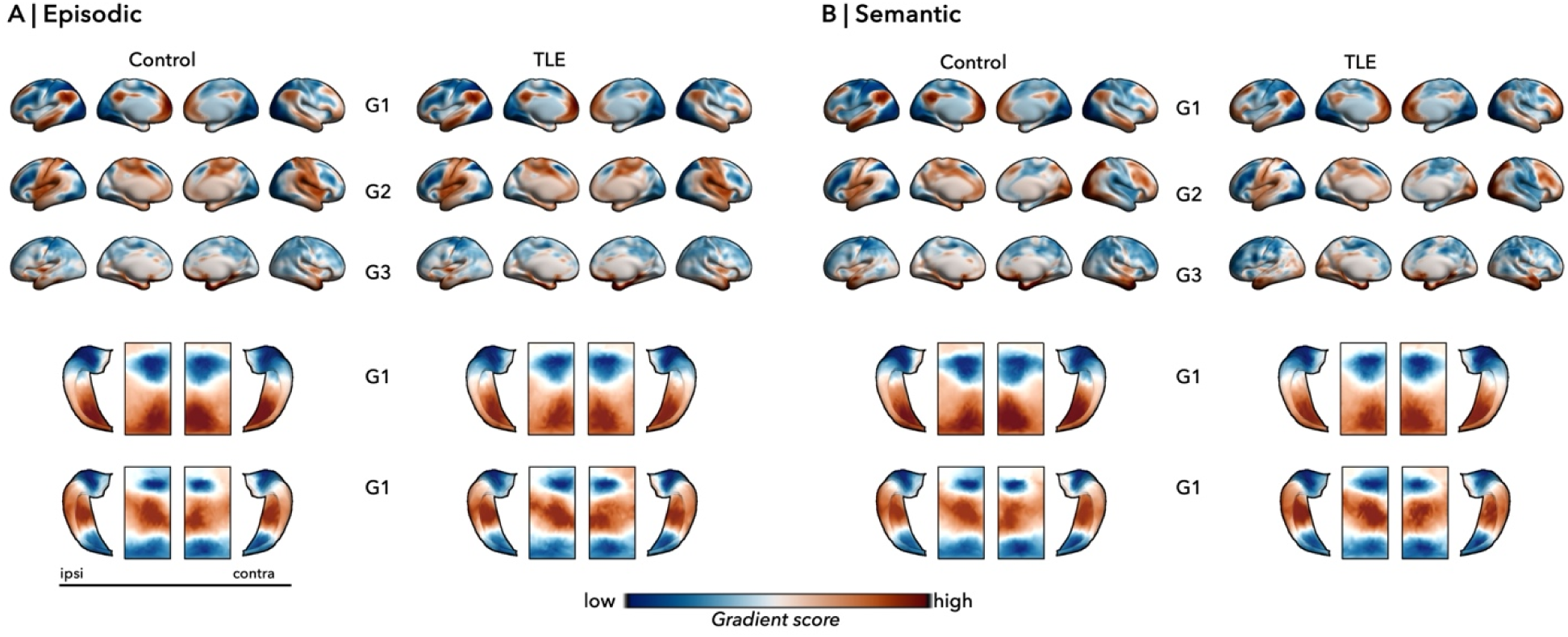
**A,B.** Episodic and semantic neocortical and hippocampal connectivity gradients before alignment to normative gradients derived from resting-state fMRI. **Abbreviations**: G1 =Gradient 1,G2= Gradient 2,G3 = Gradient 3, Ipsi= ipsilateral, Contra=contralateral, TLE=temporal lobe epilepsy

**Figure S2.**
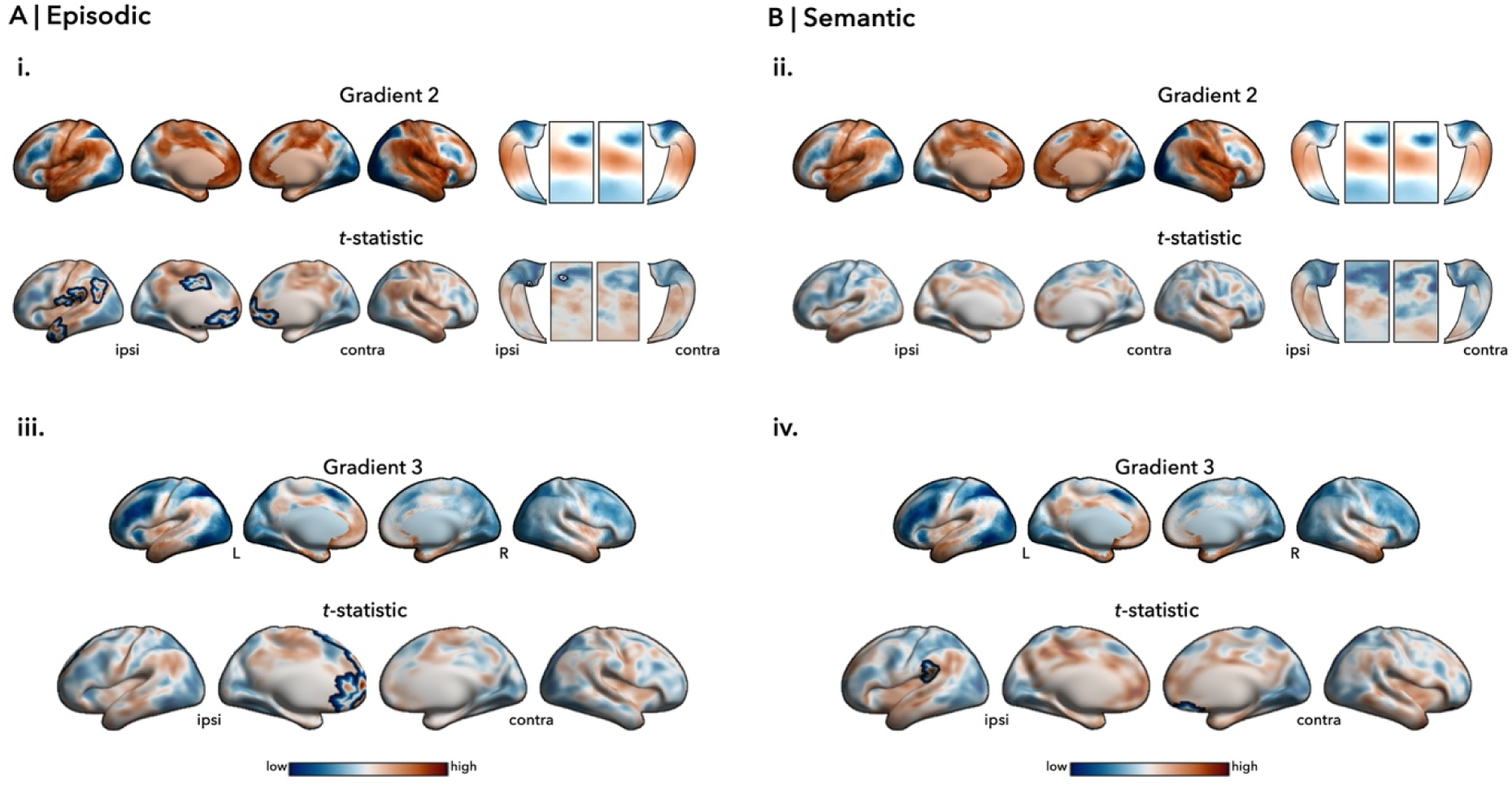
**(i,ii)**. Episodic/Semantic G2 describing a visual-to-somatomotor topography and explained 10/9% of variance. No G2 contractions were observed in semantic memory, contrasting the TLE-related reorganization observed in the anterior temporal, insular, parietal, and bilateral frontal regions for episodic memory. Significant *p*-values were highlighted after correcting for multiple comparisons (p_FWE_ < 0.05, *black outlines*). In both memory states, hippocampal G2 reflected a proximal-distal representation and explained 10/10% of functional connectome variance. While no significant differences were observed in semantic hippocampal G2, a distinct reorganization in the anterior hippocampal episodic G2 was observed. **(iii, iv).** Episodic/Semantic G3, dominated by networks involved in the declarative memory tasks explained 8% of the total episodic and semantic variance. G3 suppression in TLE for episodic memory state was recorded in prefrontal cortices, ipsilateral to seizure focus. **Abbreviations**: G2= Gradient 2, G3 = Gradient 3, Ipsi= ipsilateral, Contra=contralateral, L=left, R=right.

**Figure S3.**
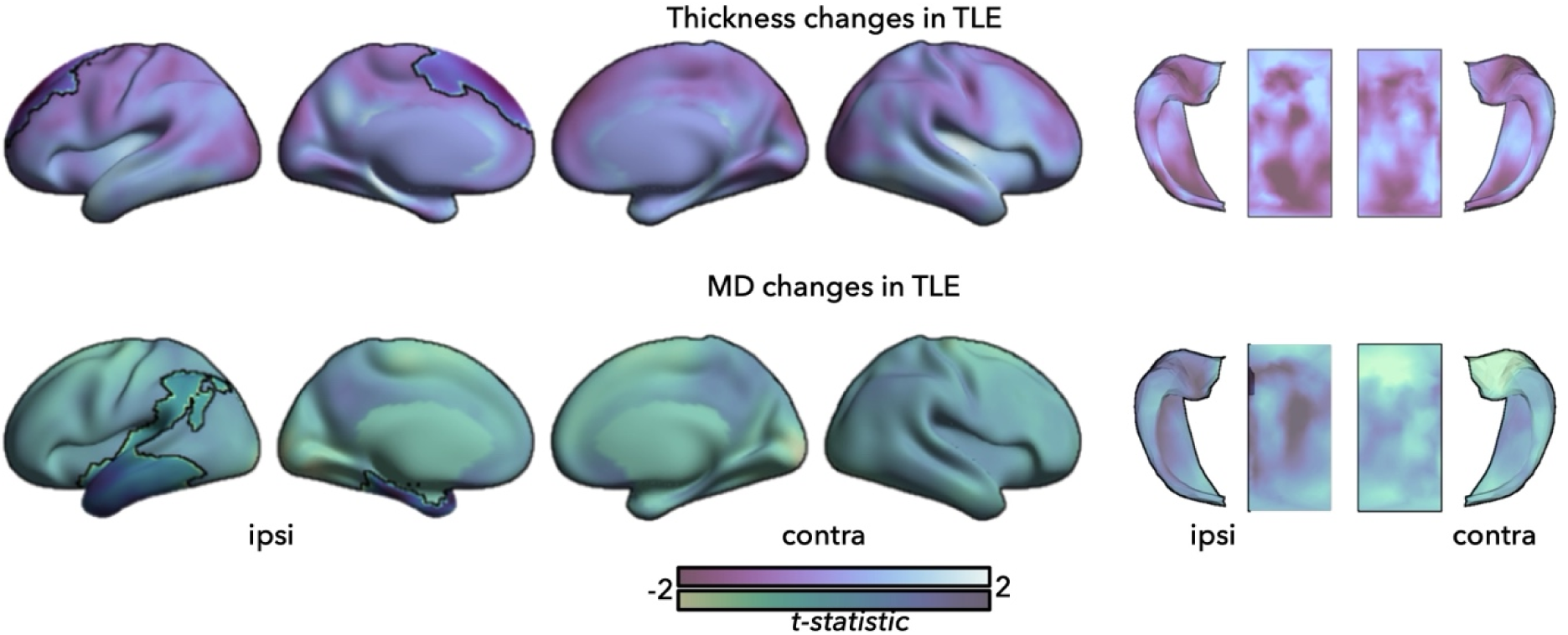
Morphology and Microstructure. Cortex-wide and subregional hippocampal univariate differences in thickness and mean diffusivity (MD) between healthy and TLE patients. Significant *p*-values were highlighted after correcting for multiple comparisons (p_FWE_ < 0.05, *black outlines*). **Abbreviations:** TLE=temporal lobe epilepsy, HC= healthy controls, Ipsi= ipsilateral, Contra=contralateral

**Figure S4.**
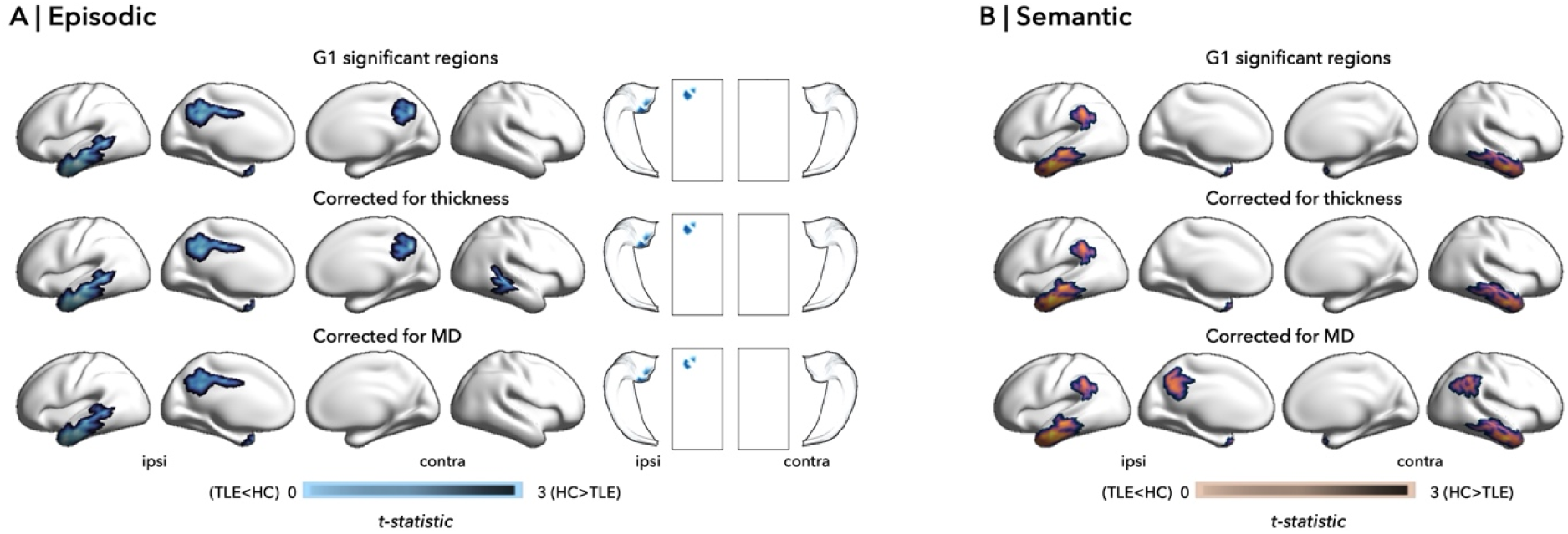
Morphology, microstructure, and gradients. Cortex-wide and subregional hippocampal univariate differences in Episodic G1 **(A)** and Semantic G1 **(B)**, after controlling for thickness and mean diffusivity (MD) between healthy and TLE patients (p_FWE_ < 0.05). At the neocortical level, thickness did not have a significant impact in both memory states, while MD significantly weakened episodic G1 and strengthened semantic G1 differences. At the hippocampal level, both morphology and microstructural properties did have a significant impact on episodic G1 differences. **Abbreviations:** TLE=temporal lobe epilepsy, HC= healthy controls, Ipsi= ipsilateral, Contra=contralateral

**Figure S5.**
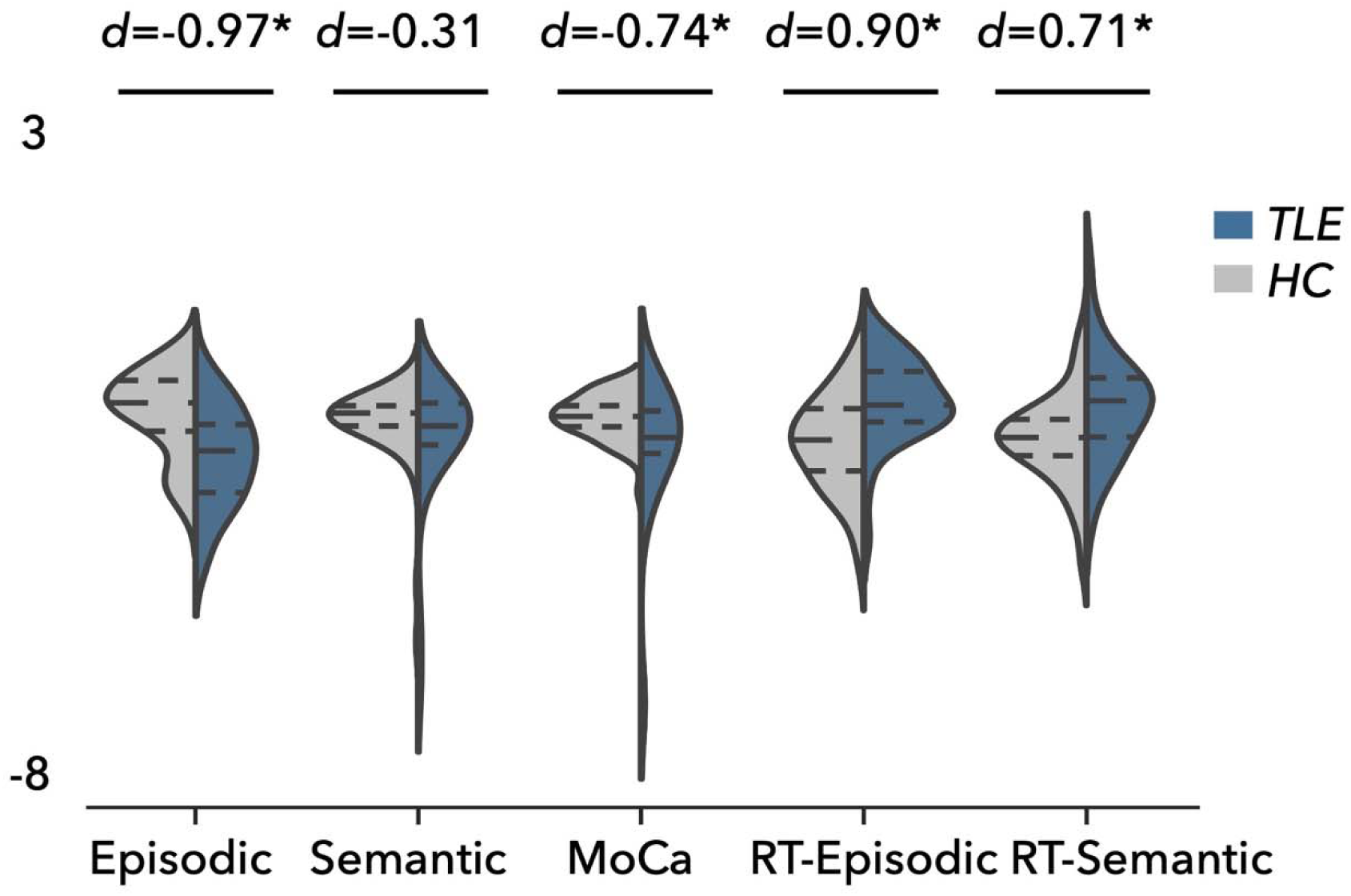
Differences in total episodic, semantic task performance, MoCA scores, and reaction times. **Abbreviations**: RT=reaction times, TLE=temporal lobe epilepsy, HC=healthy controls.

**Figure S6.**
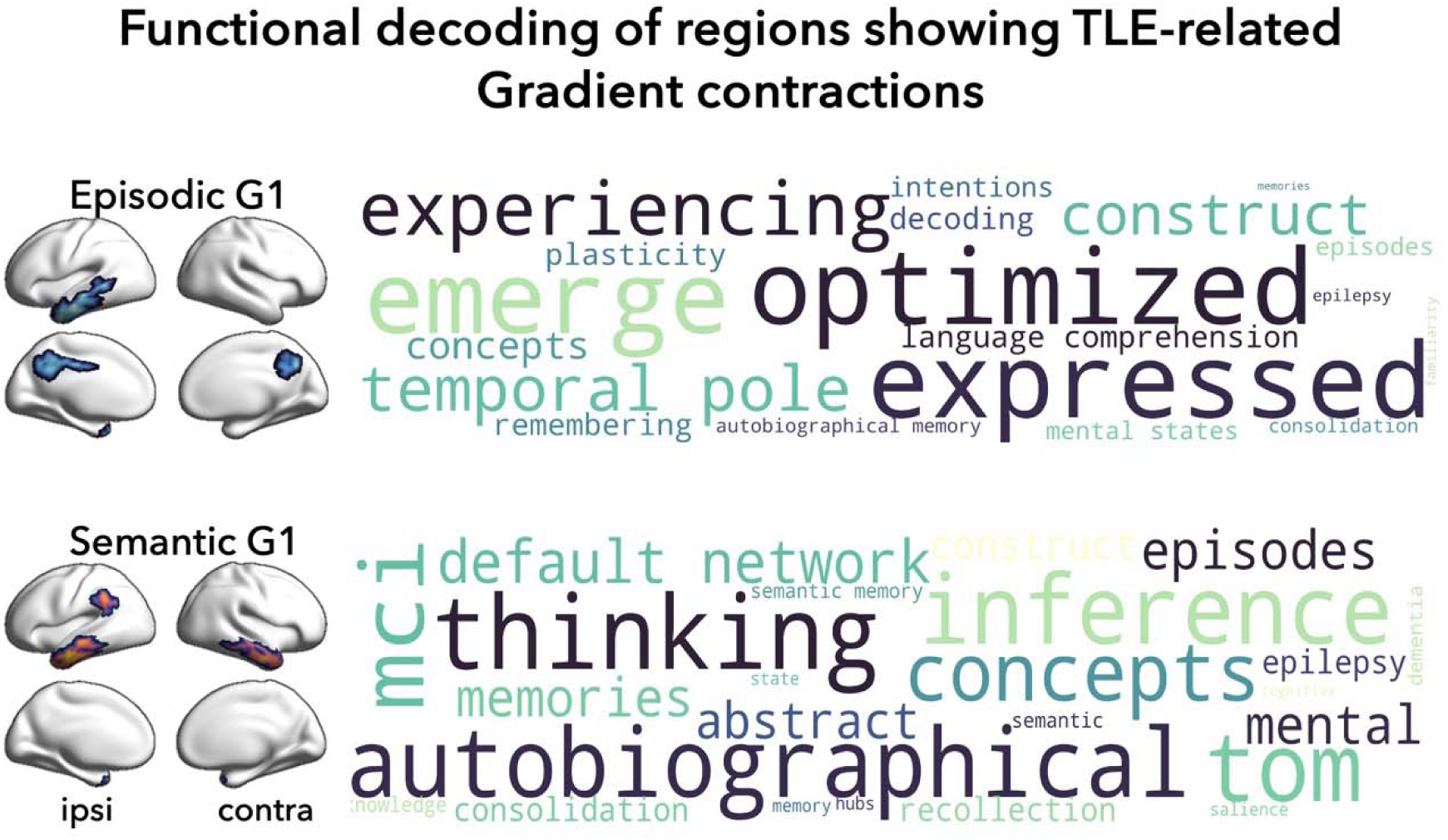
The associated cognitive terms were extracted using the *Neurosynth* decoder function and represented in a word-cloud plot. The size of the fonts used for each term reflects the correlation between the FDR-corrected maps and the *Neurosynth*-generated meta-analytic maps.

